# ADAM17 regulates hepatic DNA damage repair and tumour formation

**DOI:** 10.1101/2024.07.30.605791

**Authors:** Manuel Reichert, Birgit Halwachs, Julia Bolik, Anke Senftleben, Freia Krause, Luisa Conrady, Renate Bauer, Jacob Rachmilewitz, Jutta Horejs-Hoeck, Stefan Rose-John, Dirk Schmidt-Arras

**Affiliations:** Institute of Biochemistry, Christian-Albrechts-University Kiel, Germany; Department of Biosciences and Medical Biology, Paris-Lodron-University Salzburg, Austria; Center for Tumor Biology and Immunology, Paris-Lodron-University Salzburg, Cancer Cluster Salzburg, Austria; Goldyne Savad Institute of Gene Therapy, Haddassah Medical Center, Faculty of Medicine, Hebrew University of Jerusalem, Israel

**Keywords:** DNA damage repair, HCC, ADAM17, HB-EGF, EGFR

## Abstract

**Background:** Hepatocellular carcinoma (HCC) is one of the leading causes of cancer deaths worldwide. Still, therapy options for this tumour entity are limited and novel therapeutic options are highly sought after. Genomic instability of hepatocytes promotes oncogenic transformation and underlies the regulation of micro-environmental signalling cues. The membrane-bound a disintegrin and metalloprotease (ADAM) 17 is a major regulator of micro-environmental signals through the proteolytic release of paracrine factors. However, its role in hepatic DNA damage repair and its contribution to hepatic tumourigenesis is still unclear.

**Methods:** We investigated the effect of ADAM17 on diethylnitrosamine (DEN)-induced acute DNA damage and subsequent DNA damage repair by utilizing mice with ubiquitous or myeloid-specific genetic deficiency in ADAM17. DNA double strand breaks and inflammation were investigated by immunofluorescence of liver tissue sections. tumourigenesis in mice with myeloid-specific ADAM17-deficiency was investigated in a chemically induced hepatocarcinogenesis model.

**Results:** ADAM17 on myeloid cells, in particular Kupffer cells is essentially involved in the non-cell autonomous regulation of DNA damage repair in hepatocytes. Parenchymal ADAM17 regulates hepatocyte fate and recruitment of infiltrating myeloid cells. Furthermore, myeloid ADAM17 promotes hepatic tumour initiation and correlates with poor prognosis in human HCC.

**Conclusions:** We identified ADAM17, in particular on myeloid cells as an essential driver of hepatic tumourigenesis and as a potential novel drug target for the treatment of hepatic malignancies.

## Introduction

Cells are constantly exposed to genotoxic stress derived from metabolic or environmental sources. Efficient repair of DNA double-strand breaks (DSBs) is key to maintaining genomic stability. The DNA damage response (DDR) is a complex molecular mechanism that relies largely on protein networks in the nucleus to sense DNA damage and to recruit DNA repair enzymes to sites of DNA damage [1]. This includes the recruitment and phosphorylation of the alternative histone H2Ax (γH2Ax) to sites of DNA DSBs. While DDR is a cell-autonomous process, DNA DSB rejoining can be accelerated by the activation of the receptor tyrosine kinase epidermal growth factor receptor (EGFR) [2]. EGFR activation requires ligand binding and subsequent receptor dimerisation. Seven different EGFR ligands have been identified, which all are synthesised as trans-membrane precursors that require limited proteolysis for release [3]. Secretion of Proheparin-binding EGF-like growth factor (HB-EGF) from macrophages has been shown to enhance DNA damage repair in neighbouring epithelial cells [2]. Furthermore, cells exposed to DNA damage are sensitised to EGFR signalling via downregulation of Mig-6 [4], a negative regulator of EGFR. Interestingly, the DDR-promoting effect of EGFR is attenuated by the concomitant activation of TNFR1 by TNF, which is secreted under chronic inflammatory settings such as ageing or obesity [5].

Activation of the EGFR in myeloid cells has been shown to promote tumour formation of both, colon and hepatocellular carcinoma. Genetic ablation of EGFR in myeloid but not in epithelial cells reduced the number of tumour nodules in mouse models of colon cancer and HCC. Mechanistically, EGFR activation in myeloid cells is needed for the transcriptional up-regulation of the pro-inflammatory cytokine Interleukin 6 (IL-6) [6, 7]. Consequently, IL-6 in complex with the soluble IL-6 receptor (sIL-6RA) promotes tumourigenesis of epithelial cells in mouse models of HCC and colon cancer [8, 9]. Binding of IL-6 to its cognate trans-membrane receptor IL-6RA results in the engagement of a GP130 dimer which constitutes the signal transducing subunit of the IL-6 receptor complex. While GP130 is ubiquitously expressed, the expression of IL-6RA is restricted to certain epithelial cells and leukocytes [10]. However, IL-6 is also able to bind to sIL-6RA, which in turn can bind and activate GP130 on target cells that do not express the membrane-bound IL-6RA. This signalling mode was termed “IL-6 trans-signalling” [11]. sIL-6RA was shown to be primarily generated via limited proteolysis catalysed by trans-membrane proteases [10]. Interestingly, the tumour promoting effect of IL-6 in colon and hepatocellular cancer is mainly mediated by IL-6 trans-signalling [8, 9].

Proteolytic release of ectodomains, a process termed “ectodomain shedding”, is an irreversible post-translational mechanism that regulates protein function, intracellular signalling and the provision of signals to neighbouring cells. Members of the a disintegrin and metalloprotease (ADAM) family are membrane-bound proteases that are important mediators of ectodomain shedding. The family member ADAM17 was shown to be involved in the shedding of a plethora of transmembrane proteins, including growth factors, cytokines and receptor molecules [12]. For examples, ADAM17 has been shown shown to release several of the membrane-bound EGFR ligands including transforming growth factor α (TGFα), HB-EGF, amphiregulin (AREG) and epiregulin (EREG) [13]. Furthermore, the proteolytic generation of sIL-6RA from leukocytes was shown to be mainly mediated by ADAM17 [14].

Genetic deficiency of ADAM17 in mice is embryonic lethal [15]. Also in humans, loss of ADAM17 catalytic activity is associated with severe disease [16, 17]. Both observations are mainly linked to impaired release of EGFR ligands in the absence of ADAM17 activity. We previously generated viable ADAM17 hypomorphic (ADAM17^ex/ex^) mice with about 5% residual ADAM17 expression in all tissues. These mice are viable and allow to study the role of ADAM17 *in vivo* [18]. Furthermore, mice with a conditional allele of ADAM17 allow to study the impact of cell type-specific genetic deficiency of ADAM17 [13].

While there is experimental evidence that substrates of ADAM17 are involved in hepatic tumourigenesis [19], little is known how ADAM17 impacts DNA damage repair and tumourigenesis in the liver. In the present study, we utilize different mouse strains with genetic deficiency in ADAM17 and analyse the impact of ADAM17 on DNA damage repair and hepatic tumourigenesis. We demonstrate that both, hepatocyte and myeloid ADAM17 are involved in DNA damage response and survival of hepatocytes after an acute DNA damage. We propose a dual role of myeloid ADAM17 in the control of genotoxic stress in the liver: on one hand, it enhances DNA damage repair in hepatocytes but on the other hand it also promotes hepatic tumourigenesis, most likely by enhancing survival of pre-neoplastic hepatocytes. We therefore identified a novel role of ADAM17 during hepatic tumourigenesis.

## Material and Methods

### Materials

Diethylnitrosamine (DEN, #N0258), 1,4-Bis-[2-(3,5-dichloropyridyloxy)]benzene, 3,3’,5,5’-Tetrachloro-1,4-bis(pyridyloxy)benzene (TCPOBOP, #T1443) and 1,1,3,3-tetramethoxypropane (TMP, #108383) were purchased from Sigma-Aldrich (Steinheim, Germany). Thiobarbituric acid (TBA, #8180) was from Merck (Darmstadt, Germany). DEN and TCPOBOP were diluted in sunflower oil.

### Animal model

Hypomorphic ADAM17 (ADAM17^ex/ex^) [20] have been described previously. LysM-Cre::ADAM17^flox/flox^ (ADAM17^ΔMC^) mice were homozygous for the floxed ADAM17 allele [13] and heterozygous for the Cre recombinase under the control of the lysozyme M [21] promoter. All mice were on a C57BL/6 background and housed under controlled conditions (specific pathogen free, 22 °C, 12-hour day-night cycle) and fed a standard laboratory chow ad libitum. Animal experiments were conducted according to national and European animal regulations and have been approved by local authorities of the Schleswig Holstein State Government (V242-30412/2016 (53-5/16)).

For the induction of acute DNA-damage, 6 week-old male mice were injected i.p. with 25 mg/kg bodyweight diethylnitrosamine. Mice were sacrificed 48h post injection for further analysis.

Hepatocarcinogenesis was induced as previously described [8]. In brief, 14 day-old male mice were injected i.p. with 25 mg/kg bodyweight diethylnitrosamine. Subsequently mice received ten consecutive biweekly injections with a dose of 3 mg/kg bodyweight TCPOBOP. Two weeks after the last injection, mice were sacrificed for further analysis.

### Immunohistochemistry and immunofluorescence

Murine tissue specimens were snap frozen in OCT TissueTek compound (Plano GmbH, Wetzlar, Germany) and cut into 8 μm-thick sections. Sections were fixed in acetone:methanol (1:1 v/v) for 2 min and subjected to immunofluorescent staining according to standard procedures. Antibody specifications are listed in Supplementary Table 2. Quantification of tumour area was performed using the software QuPath [22], version 0.5.1 on Ubuntu Linux 24.04.

### SDS-PAGE and immunoblotting

Samples were lysed in RIPA buffer (50 mM HEPES, pH 7.4, 150 mM NaCl, 1 mM EDTA, 2 mM EGTA, 0.5 % NP-40) supplied with 50 mM NaF, protease and phosphatase inhibitors. Proteins were separated by electrophoresis on 10 % SDS gels and transferred to PVDF membranes. Membranes were incubated with primary antibodies overnight at 4°C and with horseradish peroxidase-conjugated secondary antibodies at room temperature for 1h. An ECL substrate kit was used for detection (Thermo Scientific, Schwerte, Germany). Antibody specifications are listed in Supplementary Table 1.

### RNA extraction

RNA of snap-frozen whole liver tissue was extracted using TRIzol (LifeTechnologies, Darmstadt, Germany) according to the manufacturer’s instructions, followed by phenol/chloroform purification. Briefly, ca. 25 mg liver tissue was removed from the left liver lobe and homogenized in TRIzol using Precellys® homogenizer and chloroform was added. After centrifugation, the aqueous phase was precipitated with isopropyl alcohol over night at -20 °C. RNA was washed with ethanol and dissolved in DEPC water before purification with phenol/chloroform using Phase Lock Gel tubes. The purified RNA was precipitated with isopropyl alcohol and 0.3 M sodium acetate overnight at -20 °C. RNA was washed with ethanol twice and dried at room temperature before dissolving in DEPC-treated water.

### Quantitative reverse transcription with PCR (RT-PCR)

Equal amounts were used for reverse transcription. One microgram of total RNA was used for reverse transcription with oligo-(dT) 18 primers and RevertAid Reverse Transcriptase (Thermo Scientific, Schwerte, Germany) and subsequently subjected to quantitative polymerase chain reaction on a Roche Lightcycler^®^ 480 using the Roche Universal Probes and Lightcycler^®^ 480 Probes Master. Primer sequences were designed using the Roche Universal Probe Library Assay Design Center and are detailed in Supplementary Table 2. Equal amounts of each sample cDNA were mixed with the respective primer and 20x Assay Universal Probe to a total volume of 10 µl per well. All samples were run in duplicates. C_T_ values were calculated using the LightCycler^®^ 480 Software 1.5.0. Respective gene expression levels were normalised to the expression of GAPDH or tubulin (ΔC_T_) and assessed by efficiency corrected calculation using a diluted pool of all cDNA samples as previously described [8, 23]. Relative gene expressions (E^-ΔCt^) are shown as fold changes relative to normalised levels in the corresponding control mice.

### Determination of lipid peroxidation

Thiobarbituric acid reactive substances (TBARS) were used as an index of lipid peroxidation and oxidative stress. In brief, proteins were precipitated from 1 mg of liver tissue lysate using 2 volumes of trichloracetic acid. The supernatant was mixed with an equal volume of thiobarbituric acid (TBA) and incubated at 100 °C for 10 min. After cooling down to room temperature, the formation of the TBA-malon dialdehyde adduct was determined photometrically at 532 nm in a Tecan plate reader. 1,1,3,3-tetramethoxypropane (TMP) was used as a standard.

### Analysis of single cell RNA sequencing data

The tabula sapiens [24] liver dataset was downloaded from figshare (TS_liver.h5ad). The human HCC was downloaded from the gene expression omnibus (GEO, accession code GSE149614). Four samples from two patients (tumour versus normal per patient) were included in the analysis (GSM4505960, GSM4505961, GSM4505963 and GSM4505964). Dimensionality reduction and gene expression analysis was generally conducted using functions implemented in the monocle3 [25] package (version 1.3.7). In combination with normalization and scaling of raw counts, dimensionality reduction was performed via principal component analysis (PCA) using the function preprocess_cds(). Next, batch correction was conducted on principal components using the align_cds() function based on the batchelor package2 [26]. Finally, the function reduce_dimension() was applied for the generation of UMAP projections. Gene expression plots were created using the function plot_genes_by_group() where groups were based on the annotations provided by the original creators of the datasets. Analysis of ligand-receptor interactions was performed via Cellchat (version 2.1.2 [27]). The analysis was conducted on normalized counts retrieved from the monocle3 cell_data_set via the function normalized_counts(). Human ligand-receptor pairs were retreived from the CellChatDB database. All data analyses were performed using R version 4.4.1.

### Data analysis and statistics

Data are represented as mean ± s.e.m. All experimental data were checked for consistency and normal distribution by using Kolmogorov-Smirnov or Shapiro-Wilk tests. Comparisons between two groups were performed by applying Student’s *t* test with and without the assumption of variance homogeneity. Wilcoxon signed rank sum test was used for paired observations. Mann-Whitney *U* test was used to compare two independent groups nonparametrically. All reported tests were two sided and two sided P<0.05 was considered statistically significant. All statistical analyses were performed using R 4.4.1 with RStudio 2024.0.2 on Ubuntu Linux 24.04 or GraphPad Prism software version 5 and 7.

## Results

### A disintegrin and metalloprotease (ADAM) 17 controls hepatic DNA damage

In order to assess the impact of ADAM17 on hepatic DNA damage control, we applied an acute DNA damage model to ADAM17^ex/ex^ hypomorphic mice, which display an ubiquitous 95% reduction in ADAM17 expression [20]. To this end, mice were injected i.p. with 25 mg/kg bodyweight (BW) diethylnitrosamine (DEN) (Fig. 1 A) which is metabolized exclusively in pericentral hepatocytes in a cytochrome P450 (CYP450)-dependent manner, resulting in DNA double strand breaks in these cells [8]. In accordance with a DEN-induced oxidative liver tissue damage we observed an increase in lipid oxidation upon acute DNA damage as assessed by malondialdehyde (MDA) levels in total liver extracts. Consequently, serum levels of alanine aminotransferase (ALT), which is released from damaged hepatocytes, were elevated (Fig. 1 B). While we did not observe a difference in the number of cells that acquired DNA damage (Fig. 1 C), the number of DNA double strand breaks per cell was significantly increased in ADAM17-deficient hepatocytes as assessed by the incorporation of the phosphorylated alternative γH2Ax histone that marks DNA double strand breaks (Fig. 1 C). As a result, we observed increased induction of apoptosis in ADAM17-deficient hepatocytes reflected by enhanced caspase 3 cleavage (Fig. 1 D+E) and an increase in DNA fragmentation as assessed by TUNEL staining (Fig. 1 E). These data indicate that while oxidative stress was similar in both groups, ubiquitous loss of ADAM17 resulted in increased genomic instability and a decrease in hepatocyte survival. However, this was not linked to altered expression of genes involved in DNA damage repair or cell cycle control (Fig. S1 A+C). Increased apoptosis was associated with enhanced phosphorylation and protein stabilisation of the tumour suppressor p53 (Fig. 1 F). While we did not detect enhanced activation of the DNA damage-sensing kinases ataxia telangiectasia mutated (ATM) and ataxia telangiectasia and Rad3-related protein (ATR), we observed slightly enhanced phosphorylation of the stress-activated MAP kinase p38 in the absence of ADAM17, which might account for the phosphorylation of p53 (Fig. 1 F).

**Figure 1.**
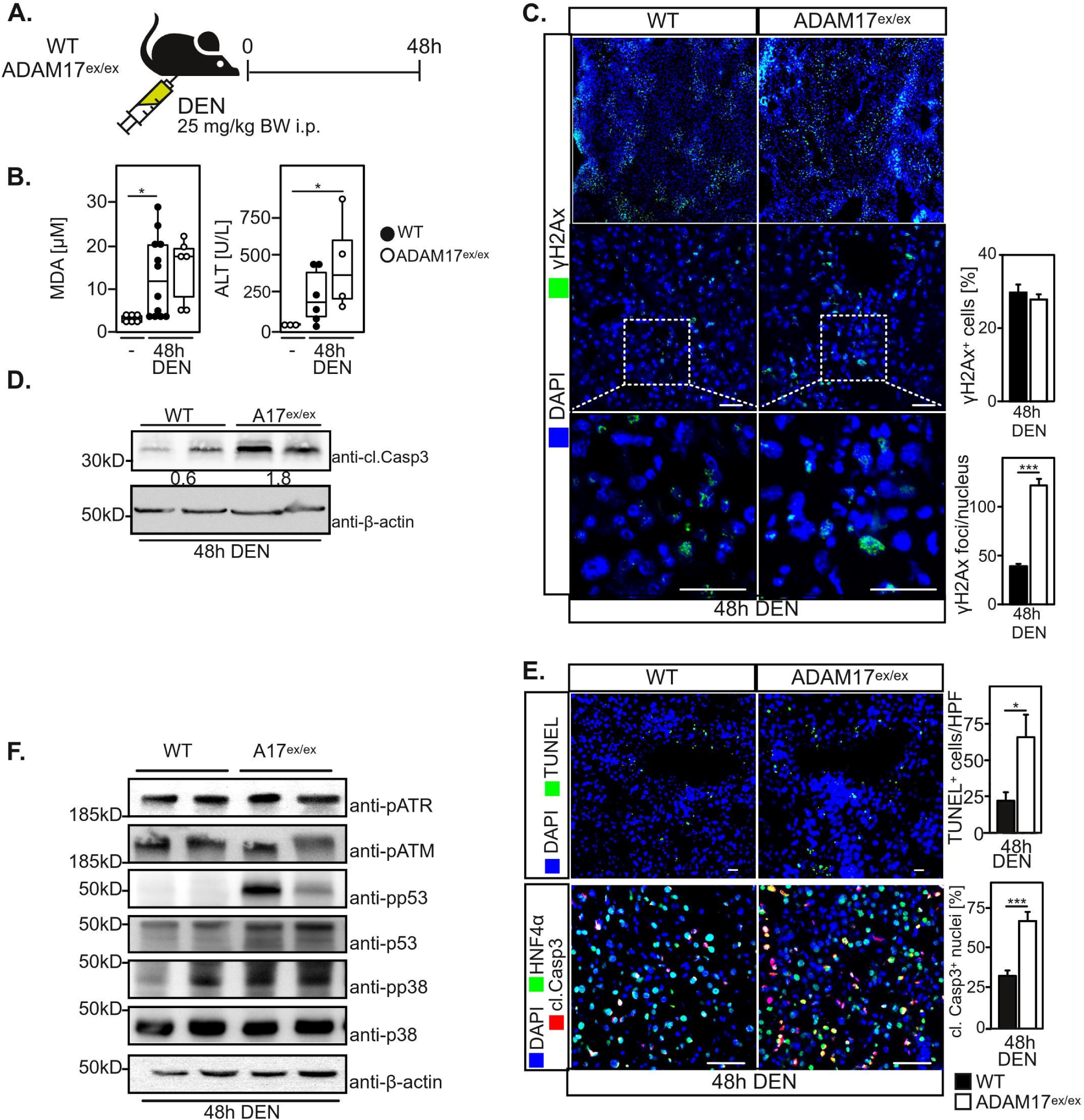
ADAM17 regulates acute DNA damage and hepatocyte survival in the liver. **A.** Experimental outline. Control (ctrl) or ADAM17-hypomorphic (ADAM17^ex/ex^) mice were i.p. injected with 25 mg/kg bodyweight (BW) diethylnitrosamine (DEN) and sacrificed after 48 hours. **B.** Malondialdehyde (MDA) levels as measure of lipid oxidation are increased upon acute DNA damage but independent of genetic ADAM17-deficiency. Serum levels of alanine-aminotransferase (ALT) are slightly enhanced in the absence of ADAM17. n=3-6 mice/group, \**P*<0.05 by unpaired two-tailed Student’s *t* test (MDA levels) or Wilcoxon rank sum test (ALT levels). **C.** Number of DNA double strand breaks per hepatocyte is enhanced in the absence of ADAM17. While the total number of γH2Ax^+^ cells is equal, the number of γH2Ax per nucleus is enhanced in the absence of ADAM17 as assessed by immunofluorescence. Scale bar represents 50 µm. n=4 mice/group, \*\*\**P*<0.001 by two-sided Wilcoxon rank sum test. **D.** Apoptosis is enhanced in the liver as assessed by cleaved caspase 3 (cl. Casp3) immunoblotting of total liver lysates. **E.** Apoptosis of hepatocytes is enhanced in the absence of ADAM17 as assessed by terminal UDP end nick labeling (TUNEL) and cleaved caspase 3 immunofluorescence of liver tissue sections. n=3 mice/group, \**P*<0.05, \*\**P*<0.01 by unpaired two-tailed Student’s *t* (cl. Casp3) or Wilcoxon rank sum (TUNEL) test. **F.** p53 is stabilised in the absence of ADAM17 as assessed by immunoblotting. Data represent mean ± standard error of mean (SEM)

Taken together, our data indicate that ADAM17 controls the persistence of hepatocytes to DNA damage.

### ADAM17 impairs myeloid cell recruitment to the liver

We next assessed if ADAM17 is involved in the induction of inflammation upon hepatic DNA damage. Immunofluorescent staining of liver tissue sections revealed significantly increased numbers of F4/80^+^ myeloid cells and an enhanced clustering of these cells in hepatic areas of DNA damage in hypomorphic ADAM17^ex/ex^ mice (Fig. 2A). In order to differentiate if this was due to enhanced proliferation of tissue-resident Kupffer cells or increased infiltration of bone-marrow-derived myeloid cells, we stained liver tissue sections with CLEC4F and CD11b. While we observed only a slight increase in CLEC4F-positive tissue-resident Kupffer cells (Fig. 2 B), the number of CD11b^+^ infiltrating myeloid cells was markedly enhanced (Fig. 2 C). We therefore concluded that ADAM17 regulates in particular infiltration of bone marrow-derived myeloid cells to sites of DNA damage in the liver. The cell adhesion molecule ICAM1 is expressed on liver sinusoidal endothelial cells and confers infiltration of myeloid cells into the liver [28]. We therefore analysed expression and localisation of ICAM1 in the liver upon acute DNA damage. Expression of *Icam1* was increased upon acute DNA damage but independent of genetic ADAM17-deficiency (Fig. 2 C). ICAM1 localised pericentrally but did not display alterations in the absence of ADAM17 (Fig. 2 D). We therefore analysed the expression of the myeloid-attracting chemokines CCL2 and CCL3 [28]. Expression of both, *Ccl2* and *Ccl3* was increased upon DNA damage. *Ccl2* expression was more strongly elevated in the liver of mice with ubiquitous ADAM17-deficiency (Fig. 2 E). In line with enhanced recruitment and activation, expression of the inflammatory cytokine genes *Tnfa* and *Il1b* was increased in the liver upon acute DNA damage. Expression of *Tnfa* was significantly increased in ADAM17-deficient mice as compared to control animals (Fig. 2 F).

**Figure 2.**
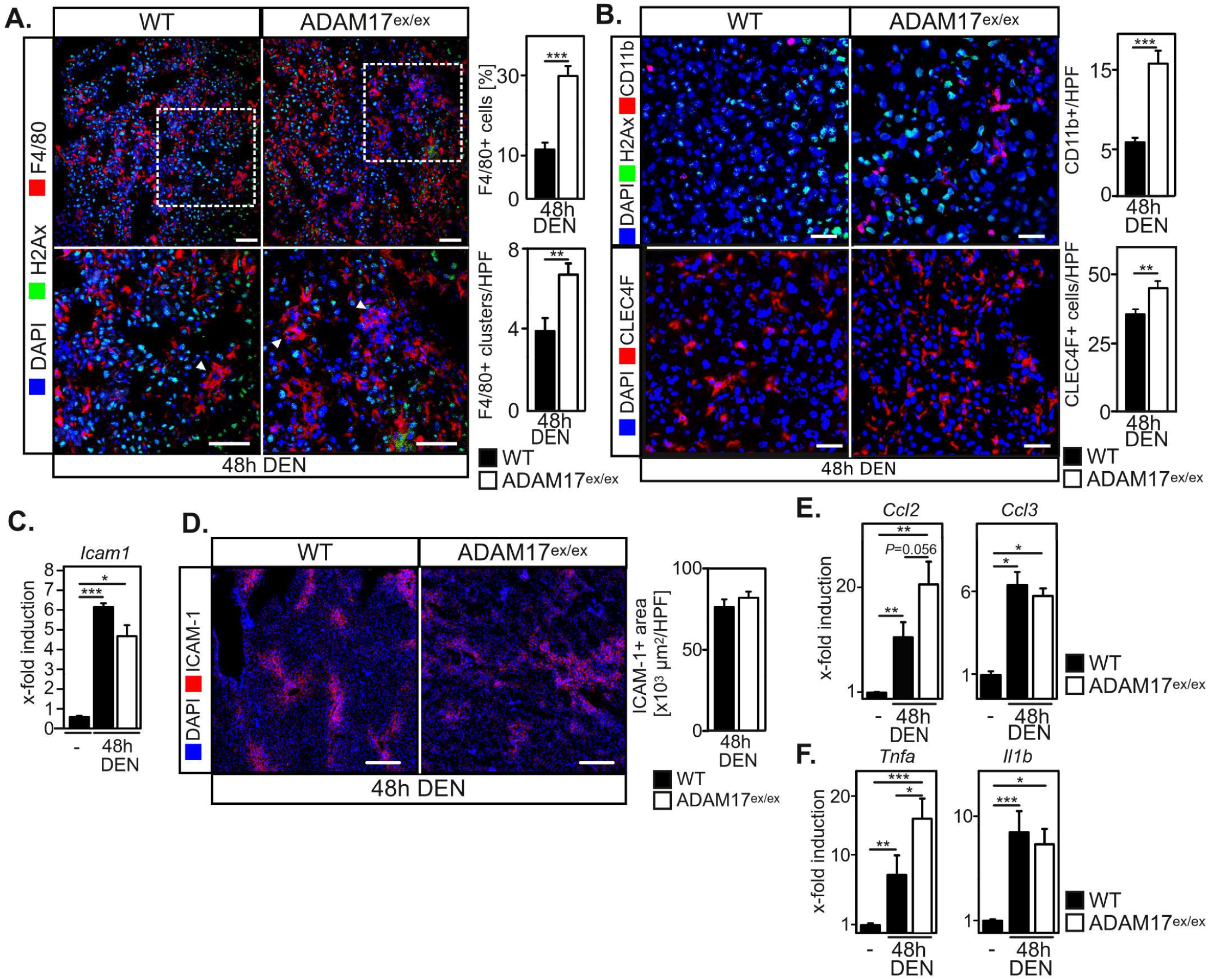
ADAM17 regulates myeloid cell recruitment to sites of acute hepatic DNA damage. **A.** The number of F4/80^+^ myeloid cells and their aggregation into clusters is strongly enhanced in ADAM17-deficient mice upon acute DNA damage as assessed by immunofluorescence of liver tissue sections. Scale bar indicates 30 µm. n=4 mice/group, \*\**P*<0.01, \*\*\**P*<0.001 by unpaired two-tailed Student’s *t* test. **B.** The number of CD11b^+^ infiltrating macrophages and of CLEC4F^+^ tissue resident Kupffer cells in the liver is enhanced in ADAM17-deficient mice upon acute DNA damage as assessed by immunofluorescence of liver tissue sections. Scale bar indicates 30 µm. n=3-4 mice/group, \*\*\**P*<0.001 by unpaired two-tailed Student’s *t* test. **C.** Expression of the cell adhesion molecule ICAM1 is enhanced in the liver upon acute DNA damage but not dependent on the presence of ADAM17, as assessed by qRT-PCR of total liver RNA. n=3 mice/group, \*\*\**P*<0.001 by unpaired two-tailed Student’s *t* test. **D.** Spatial expression of ICAM1 is not altered between wildtype and ADAM17-deficient mice upon acute DNA damage in the liver as assessed by immunofluorescence of liver tissue sections. n=4 mice/group, unpaired two-tailed Student’s *t* test. **E.** Expression of myeloid chemokines *Ccl2* and *Ccl3* are enhanced upon hepatic acute DNA damage as assessed by qRT-PCR of total liver RNA. Expression of *Ccl2* is enhanced in the absence of ADAM17. n=3 mice/group, \**P*<0.05, \*\**P*<0.01 by Wilcoxon signed rank sum test (Ccl2) and unpaired two-tailed Student’s *t* test (Ccl3). **F.** Expression of the myeloid cytokines TNF and IL1β is increased upon acute hepatic DNA damage as assessed by qRT-PCR of total liver RNA. TNF but not IL1β is enhanced in the absence of ADAM17. Data represent mean ± standard error of mean (SEM).

Taken together, these data indicate that loss of ADAM17 enhances myeloid cell recruitment to the liver via enhanced secretion of CCL2.

### Myeloid ADAM17 controls hepatic DNA damage repair in a non-cell autonomous manner

Myeloid cells were shown to promote DNA damage repair in a non-cell autonomous way via secretion of Proheparin-binding EGF-like growth factor (HB-EGF) and activation of EGF receptor (EGFR) signalling in hepatocytes [2]. Furthermore, IL-1β, a cytokine that is induced upon acute hepatic damage, has been shown to up-regulate *Adam17* expression in Kupffer cells [6]. We therefore sought to investigate the role of ADAM17 on myeloid cells for DNA damage repair, particularly as ADAM17 was identified as sheddase for several EGFR ligands including HB-EGF [29, 30].

We generated mice with myeloid-specific ADAM17-deficiency by breeding LysM-Cre transgenic mice [21] to mice with a conditional allele of ADAM17 (ADAM17^fl/fl^) [13]. Recombination of the conditional allele was detectable in both, bone marrow-derived macrophages (BMDM), as well as in total liver genomic DNA (Fig. S2 A-D). ADAM17 protein was completely absent in BMDMs (Fig. S2 E) and reduced in total liver lysates (Fig. S2 F). We subjected these mice to acute hepatic DNA damage by injecting 25 mg/kg BW DEN i.p. and sacrificed animals after 48 hours or six days (Fig. 3 A). Induction of acute hepatic DNA damage was accompanied with an increase in lipid oxidation as assessed by total liver MDA levels. Surprisingly, MDA levels were significantly higher in mice with myeloid-specific ADAM17-deficiency as compared to control mice (Fig. 3 B) and higher than in mice with ubiquitous ADAM17-deficiency (Fig. 1 B). In these mice, we also observed increase in ALT serum levels, whereas ALT levels were not higher in mice with myeloid-specific ADAM17-deficiency as compared to control mice (Fig. 3 B). In contrast to mice with ubiquitous ADAM17-deficiency we did not observe an altered frequency of DNA double strand breaks in the absence of myeloid ADAM17 (Fig. 3 C). Furthermore, we did not detect significant alterations in cleaved caspas 3 cleavage in the absence of myeloid-specific ADAM17 (Fig. 3 D+E), suggesting the myeloid ADAM17 does not control induction of DNA double strand breaks or hepatocyte survival upon acute DNA damage.

**Figure 3.**
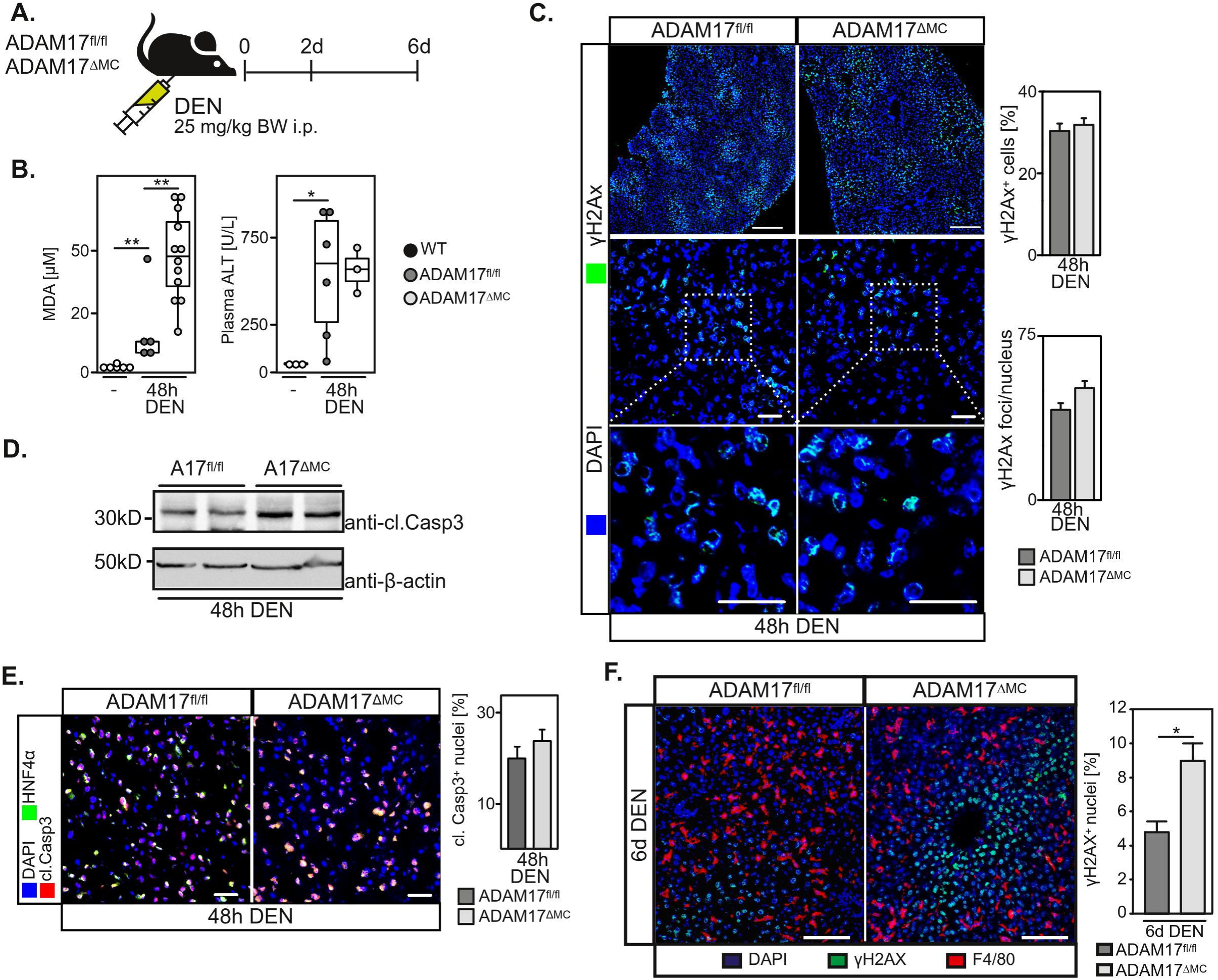
Myeloid ADAM17 is required for efficient hepatic DNA damage repair. **A.** Experimental outline. Mice with myeloid-specific ADAM17-deficiency (LysM-Cre x ADAM17^fl/fl^; ADAM17^ΔMC^) and respective control mice (ADAM17^fl/fl^) were intraperitoneally (i.p.) injected with 25 mg/kg bodyweight (BW) diethylnitrosamine (DEN) and sacrificed after 48 hours. **B.** Malondialdehyde (MDA) levels as measure of lipid oxidation are increased upon acute DNA damage but independent of genetic ADAM17-deficiency. Serum levels of alanine-aminotransferase are slightly enhanced in mice with myeloid-specific ADAM17-deficiency upon acute hepatic DNA damage. n=3-6 mice/group, \**P*<0.05, \*\**P*<0.01 by Wilcoxon signed rank sum test (ALT levels). **C.** The number of DNA double strand breaks is not altered in the absence of myeloid ADAM17 upon acute DNA damage, as assessed by γH2Ax immunofluorescence of liver tissue sections. n=4 mice/group, unpaired two-tailed Wilcoxon rank sum test. **D.** Induction of apoptosis in the liver is slightly enhanced in the absence of myeloid ADAM17 as assessed by cleaved Caspase 3 (cl. Casp3) immunoblotting of total liver lysates. **E.** Apoptosis of hepatocytes upon acute DNA damage is slightly enhanced in the absence of ADAM17 as assessed by cleaved caspase 3 immunofluorescence of liver tissue sections. n=3 mice/group, unpaired two-tailed Student’s *t* test. **F.** DNA damage repair is prolonged in the absence of myeloid ADAM17. DNA damage was induced by i.p. injection of 25 mg/kg BW DEN. Mice were sacrificed two or six days post DEN-injection and DNA damage was assessed by γH2Ax immunofluorescence of liver tissue sections. n=3 mice/group, \*\**P*<0.001 by unpaired two-tailed Student’s t test with Welch’s correction. Data represent mean ± standard error of mean (SEM).

Given the prominent role of HB-EGF in non-cell autonomous control of hepatocyte DNA damage repair [2, 5], we assessed the impact of myeloid ADAM17 on the repair of DNA double strand breaks. To this end, mice were injected with 25 mg/kg BW DEN and analysed two and six days post DEN injection (Fig. 3 A). Interestingly, six days after DEN injection, the number of γH2Ax-positive nuclei was largely reduced in control animals (Fig. 3 F), but was still significantly higher in mice with myeloid deficiency in ADAM17 (Fig. 3 F). These findings indicate that myeloid ADAM17 is essentially involved in non-cell autonomous regulation of DNA damage repair and cell fate of hepatocytes.

### Myeloid ADAM17 controls hepatic EGFR activation

We analysed recruitment of myeloid cells upon acute DNA damage in mice with myeloid-specific ADAM17-deficiency. We observed reduced pericentral expression of ICAM1 in livers of mice with myeloid-specific ADAM17-deficiency as compared to control mice (Fig. S3 A) which was not associated with altered *Icam1* expression (Fig. S3 B). In contrast, to decreased ICAM1 expression, we did neither find altered numbers of CD11b+ infiltrated myeloid cells nor altered numbers of CLEC4F+ Kupffer cells in the absence of myeloid ADAM17 (Fig. S2 C). This indicates that myeloid ADAM17 does not appear to control migration of myeloid cells to the liver upon acute damage.

We next hypothesised that myeloid ADAM17 may regulate DNA damage repair in hepatocytes via the release of EGFR ligands. We therefore assessed EGFR phosphorylation upon DNA damage in control mice and mice with ubiquitous or myeloid-specific ADAM17-deficiency. While total hepatic EGFR phosphorylation was reduced in the absence of myeloid ADAM17 (Fig. 4 A), it was nearly absent in mice with ubiquitous ADAM17-deficiency (Fig. 4 B). Interestingly, we observed the most prominent reduction in EGFR phosphorylation in CLEC4F^+^ Kupffer cells, both, in mice with ubiquitous, as well as in mice with myeloid-specific ADAM17-deficiency (Fig. 4 C+D). These data indicate that upon acute hepatic DNA damage, ADAM17 promotes autocrine EGFR activation in Kupffer cells. Previous reports have suggested that autocrine EGFR activation in myeloid cells correlates with the expression of interleukin 6 (IL-6) [6, 7]. We indeed detected impaired expression of *Il6* in mice with myeloid-specific ADAM17-deficiency (Fig. 4 E). However, while we observed an increase in plasma levels of soluble IL-6 receptor (sIL-6R) upon DEN treatment, levels were equal in both, ADAM17-deficient and control mice (Fig. 4 F). This observation was not associated with an altered expression or stability of IL-6R or the homologues protease ADAM10 (Fig. S4 A-D). These data suggest that ADAM17 is not responsible for the proteolytic release of sIL-6R upon acute DNA damage in the liver.

**Figure 4.**
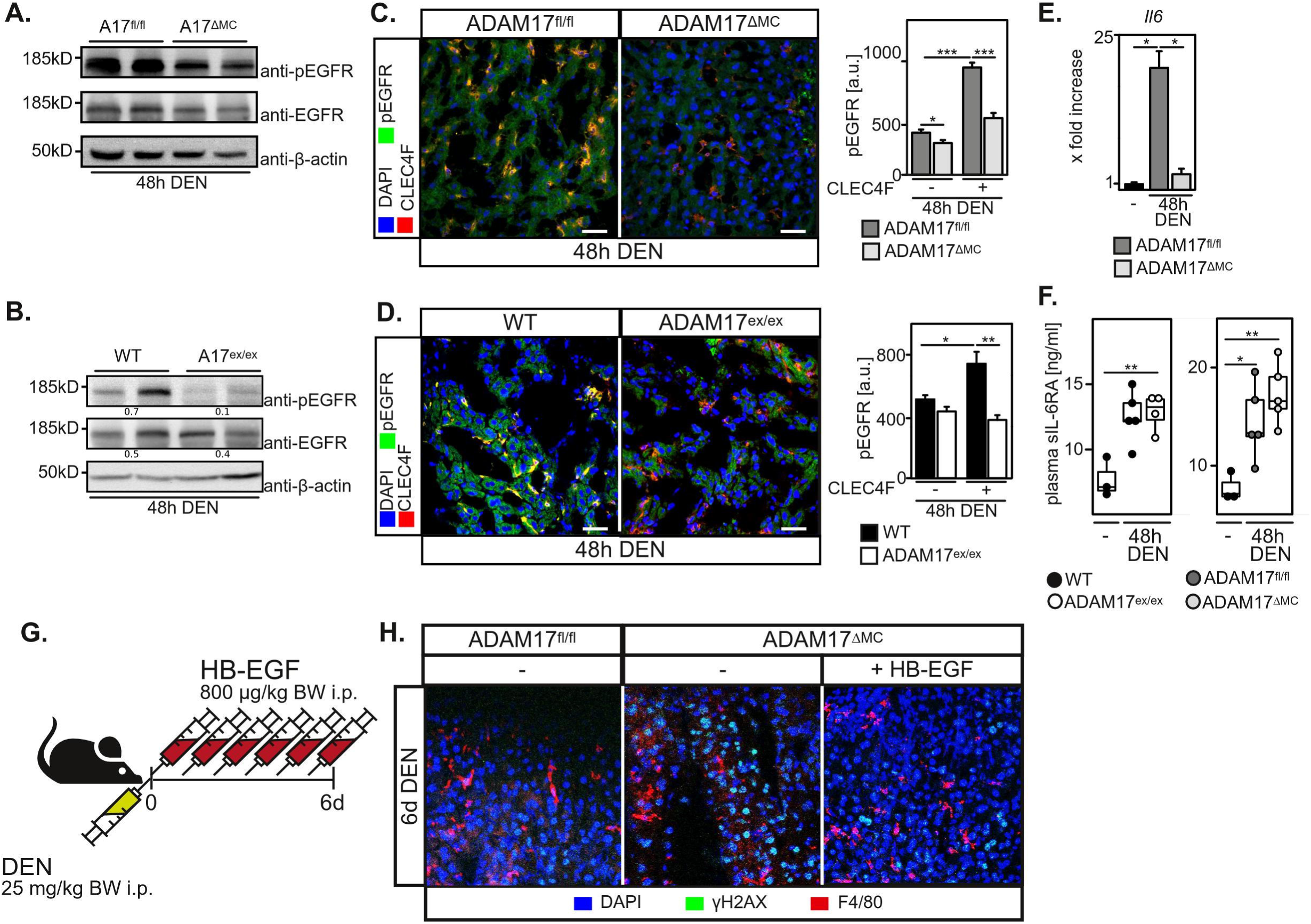
ADAM17 regulates DNA damage repair via EGF receptor activation in Kupffer cells. **A.** Hepatic EGFR phosphorylation is reduced in mice with myeloid-specific ADAM17-deficicency (ADAM17^ΔMC^) as assessed by immunoblotting of total liver lysates. **B.** Hepatic EGFR phosphorylation is completely impaired in mice with ubiquitous ADAM17-deficiency (ADAM17^ex/ex^) as assessed by immunoblotting of total liver lysates. **C.** EGFR phosphorylation is predominantly reduced in CLEC4F^+^ Kupffer cells upon acute DNA damage in the absence of myeloid ADAM17 as assessed by CLEC4F and pEGFR immunofluorescence of liver tissue sections. Scale bar indicates 30 µm. n=2-3 mice/group, \**P*<0.05, \*\*\**P*<0.001 by Wilcoxon signed rank sum test. **D.** EGFR phosphorylation is predominantly reduced in CLEC4F^+^ Kupffer cells in mice with ubiquitous ADAM17-deficiency upon acute DNA damage as asessed by CLEC4F and pEGFR immunofluorescence of liver tissue sections. Scale bar indicates 30 µm. n=3 mice/group, \**P*<0.05, \*\**P*<0.01 by Wilcoxon signed rank sum test. **E.** Induction *Il6* expression upon acute DNA damage is impaired in the absence of myeloid ADAM17, while induction of *Il6ra* expression is unaltered, as assessed by qRT-PCR of total liver RNA. n=3 mice/group, \**P*<0.05 by unpaired two-tailed Student’s *t* test. **F.** Plasma levels of soluble IL-6 receptor (sIL-6R) are significantly increased upon acute DNA damage but plasma levels are independent of genetic ADAM17 deficiency. n=3-5 mice/group, \**P*<0.05, \*\**P*<0.01, **G.** Experimental outline of H. Mice were initially injected with 25 mg/kg bodyweight (BW) followed by six daily injections with 800 µg/kg BW HB-EGF. **H.** γH2Ax immunofluorescence of liver tissue sections demonstrates enhanced DNA damage repair in HB-EGF-treated mice with myeloid-specific ADAM17-deficiency. Data represent mean ± standard error of mean (SEM).

In order to analyse, if exogenously administered HB-EGF can compensate for the loss of myeloid ADAM17, we initially injected ADAM17^ΔMC^ mice with 25 mg/kg BW DEN to induce acute DNA damage and subsequently administered 800 µg/kg BW recombinant HB-EGF daily (Fig. 4 G). Mice were sacrificed after six days. While control mice displayed sufficient repair of DNA damage after 6 days, mice with myeloid ADAM17-deficiency still displayed signs of DNA double strand breaks (Fig. 4 H). In line with a prominent role in non-cell autonomous DNA damage repair, the number of γH2Ax^+^ hepatocytes was reduced in mice that received recombinant HB-EGF (Fig. 4 H), indicating that HB-EGF-mediated EGFR activation can compensate the loss of myeloid ADAM17.

### Myeloid ADAM17 regulates hepatic tumourigenesis

A previous report suggested that myeloid EGFR activation is critical for hepatocarcinogenesis [6]. We therefore hypothesised, that ADAM17 in myeloid cells plays a major role in hepatic tumourigenesis. We therefore investigated tumour formation in mice with myeloid-specific ADAM17-deficiency using a chemically induced hepatocarcinogenesis model. Control mice and mice with myeloid ADAM17-deficiency were injected 25 mg/kg BW DEN at 14 days of age, followed by biweekly injections of the tumour promoter and constitutive androstane receptor (CAR) ligand TCPOBOP (Fig. 5 A), as previously described [8]. Relative liver and spleen weight was slightly, but non-significantly increased in myeloid ADAM17-deficient mice (Fig. 5 B). Hepatic tumour formation was evident in both, control and ADAM17-deficient animals. However, the number of hepatic tumour nodules was reduced in the absence of myeloid ADAM17 (Fig. 5 C), consistent with the observed decrease in *Il6* expression (Fig. 4 E) and a prominent role of IL-6 trans-signalling in hepatocarcinogenesis [8]. The number of tumour nodules was significantly reduced in the absence of myeloid ADAM17 (Fig. 5 C). However, we noted that the size of individual tumour nodules was slightly but not significantly increased (Fig. 5 C+D). Finally, we wanted to assess whether myeloid ADAM17 plays a comparable role in humans. Therefore, we interrogated publicly available datasets [24]. Single-cell RNA sequencing data of human liver confirmed the prominent expression of ADAM17 in liver myeloid cells and endothelial cells (Fig. 5 E). We also confirmed expression of ADAM17 substrates such as the membrane-bound EGF receptor ligands HB-EGF, AREG and EREG, as well as the IL-6RA in liver myeloid cells (Fig. 5 E). Analysis of cell-cell communication via CellChat [31] identified prominent EGF-signaling between myeloid cells (senders) and hepatocytes (receivers) (Fig. 5 F), confirming an essential role of the EGFR pathway in myeloid cell-hepatocyte cellular interactions. When analysing single-cell RNA sequencing data of human hepatocellular carcinoma (HCC) [32], we confirmed expression of ADAM17 and EGFR ligands, as well as IL-6 and IL-6R in myeloid cells in tumour tissue (Fig. 5 G). It is therefore likely that in human liver tumours a myeloid ADAM17-mediated HB-EGF-EGFR-IL-6 axis supports tumour growth. Furthermore, RNA expression levels of ADAM17 in total liver RNA of human HCC [33] inversely correlated with patients’ survival rates (Fig. 5 G). These data indicate that ADAM17 on myeloid cells plays an important role in human hepatocellular carcinoma.

**Figure 5.**
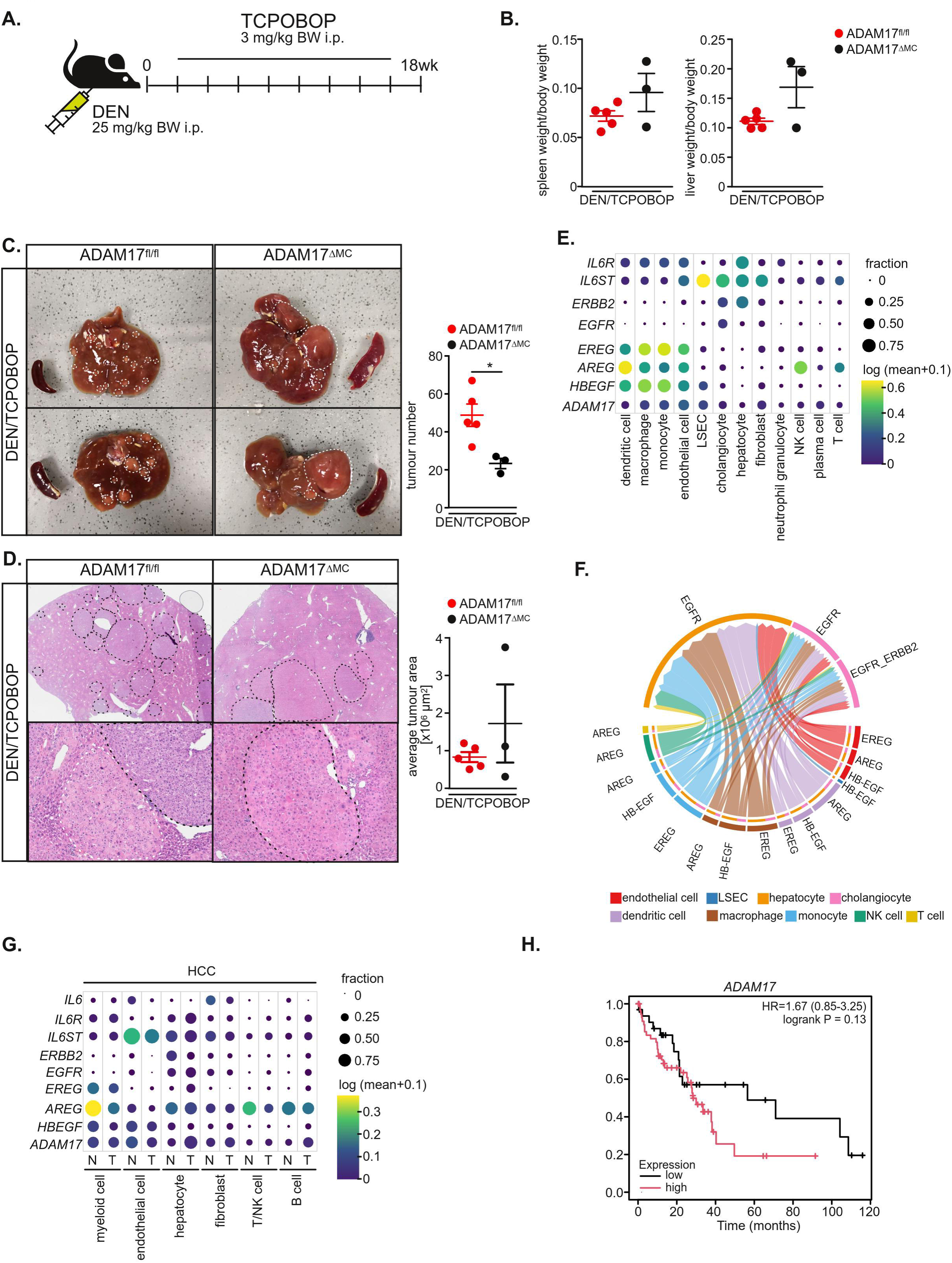
Myeloid ADAM17 affects tumour formation in the liver. **A.** Experimental outline. Control (ADAM17^fl/fl^) and ADAM17^ΔMC^ mice were i.p. injected with 25 mg/kg BW DEN at the age of 14 days. Subsequently mice were i.p. injected biweekly with the constitutive androstane (CAR) receptor ligand 1,4-Bis-[2-(3,5-dichloropyridyloxy)]benzene, 3,3’,5,5’-Tetrachloro-1,4-bis(pyridyloxy)benzene (TCPOBOP). **B.** Relative liver and spleen weight are only slightly enhanced in mice with myeloid-specific deficiency in ADAM17. **C.** The number of macroscopic visible tumour nodules is reduced in the absence of myeloid ADAM17. n=3-5 mice/group, \**P*<0.05 by Mann-Whitney *U* test. **D.** H/E staining of liver tumour sections. The frequency of tumour nodules is reduced in mice with myeloid-specific ADAM17-deficiency. The average tumour area is slightly but non-significantly increased in mice with myeloid-specific ADAM17-deficiency. **E+F.** Single cell RNA sequencing data were retrieved from public repositories [24] and expression of ADAM17 and selected ADAM17 substrates were analysed. ADAM17 and indicated EGFR ligands, as well as IL6R are predominantly expressed in myeloid cells and endothelial cells (E). Receptor-ligand interactions (F) were assessed using CellChat [31] and demonstrate that EGFR signalling on hepatocytes appears to be mainly elicited from myeloid and endothelial cells. **G.** Single cell RNA sequencing data of patients with hepatocellular carcinoma (HCC) were retrieved from public repositories [32] and expression of ADAM17 and ADAM17 substrates was analysed. ADAM17 and indicated EGFR ligands, as well as IL-6R and IL-6 are expressed in myeloid cells. N:normal tissue, T:tumour tissue **H.** Expression of ADAM17 correlates with poor prognosis in HCC patients. Patients with alcohol abuse or hepatitis infection were excluded from the data set. Data represent mean ± standard error of mean (SEM).

Taken together, we have identified myeloid ADAM17 as an essential promoter of hepatocellular carcinoma.

## Discussion

The liver is the first-pass filter of blood that drains from the intestines. Hepatocytes are therefore constantly exposed to bacterial antigens and dietary metabolites that may result in oxidative stress in the liver. Consequently, hepatocytes encounter genotoxic stress that needs to be efficiently repaired to prevent the formation of oncogenic mutations and to ensure hepatocyte homeostasis. However, during drug-induced liver injury, chronic viral hepatitis and fatty liver disease the liver is exposed to persistent oxidative stress.

DNA damage repair (DDR) is a cell-intrinsic mechanism involving multiple proteins that coordinate the repair of DNA lesions [1]. However, more recent reports have provided evidence that DDR is regulated in a non-cell autonomous manner. Kupffer cells in the liver are key regulators of hepatocyte DDR. While the release of HB-EGF promotes DDR, the release of TNF, particularly during inflamm-ageing, blocks the beneficial effects of hepatocyte EGFR activation [2, 5]. Both factors are synthesised as membrane-bound proteins and must be proteolytically released in order to act in a paracrine manner and have been shown to be processed by ADAM17 [30]. Furthermore, TNF receptor 1 has been demonstrated to be processed by ADAM17, blunting inflammatory TNF signalling but promoting cell death induction [34, 35]. We therefore hypothesised that ADAM17, in particular on myeloid cells, is critically involved in hepatocyte DDR. We observed that ubiquitous ADAM17-deficiency sensitised hepatocytes to DNA damage. The number of DNA double-strand breaks per nucleus was increased, associated with a significant upregulation of p53. Consequently, we observed a significant increase in hepatocyte apoptosis in the absence of ADAM17. Mice with myeloid-specific ADAM17-deficiency however did not show significantly altered cell death induction compared to control, suggesting that hepatocyte ADAM17 plays a cell-autonomous role in regulating acute DNA damage response. However, this was neither linked to altered expression of genes involved in the DNA double-strand break response nor in altered responses of the DNA kinases ATM or ATR. Most likely as a consequence of increased hepatocyte death, we observed increased recruitment of CD11b^+^ myeloid cells in the absence of ADAM17.

However, loss of myeloid ADAM17 delayed DDR which could be rescued by the addition of exogenous HB-EGF. In the same line, EGFR phosphorylation was reduced in the absence of myeloid ADAM17, particularly in Kupffer cells, suggesting that myeloid ADAM17 is necessary for the release of EGFR ligands and autocrine activation of myeloid EGFR. Therefore, through the release of EGFR ligands, myeloid ADAM17 plays a critical role in promoting non-cell autonomous DNA damage repair (Fig. 6 A), which is consistent with previous publications demonstrating an important role for EGFR in DDR in hepatocytes [2, 5].

**Figure 6.**
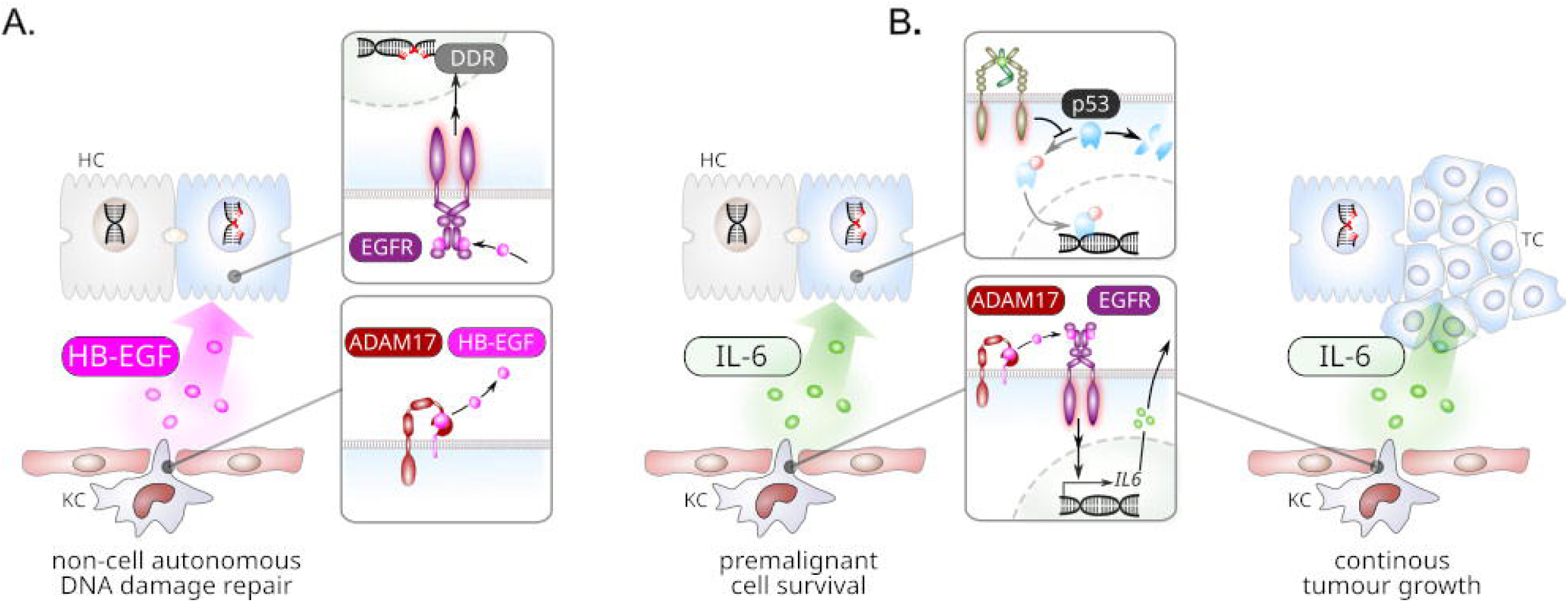
Role of myeloid ADAM17 in DNA damage control and hepato-carcinogenesis. **A.** In response to acute DNA damage, ADAM17 proteolytically releases HB-EGF from hepatic myeloid cells resulting in hepatocyte EGFR activation and enhanced cell autonomous DNA damage repair. **B.** In response to acute DNA damage, ADAM17-mediated release of EGFR ligands from liver myeloid cells induces autocrine EGFR activation and subsequent expression of IL-6. Consequently, IL-6 promotes survival of genomically unstable hepatocytes and growth of tumour lesions.

In accordance with a previous published role of myeloid EGFR for the induction of IL-6 expression [6], we observed a decrease in IL-6 in mice with myeloid ADAM17-deficiency. Therefore, myeloid ADAM17 in the liver is essential for the activation of the EGFR and subsequent induction of IL-6 and the survival of premalignant hepatocytes (Fig. 6 B). We and others have demonstrated that IL-6 [36] and IL-6 trans-signalling in particular [8] promotes hepatocarcinogenesis upon genotoxic stress. This is linked at least in part to destabilisation of p53 upon genotoxic stress and therefore to subsequent survival of genomically unstable hepatocytes. In this context we identified Kupffer cells as the major source of sIL-6R [8]. Interestingly, in our current study, we did not observe alterations in sIL-6R plasma levels upon acute DNA damage, suggesting that ADAM17 is not responsible for the proteolytic release of sIL-6R from Kupffer cells under these conditions.

Given the prominent role of myeloid ADAM17 in the DNA damage response and in particular in the upregulation of IL-6, we hypothesised that myeloid ADAM17 may be critically involved in promoting hepatocarcinogenesis. Indeed, we observed that loss of myeloid ADAM17 significantly reduced the frequency of tumour nodules, which appeared to be associated with reduced IL-6 signalling and hence reduced survival of genomically unstable hepatocytes. Therefore, myeloid ADAM17 in the liver promotes hepatocarcinogenesis (Fig. 6 B). We noted that the size of tumour nodules appeared to be slightly larger in the absence of myeloid ADAM17. Interestingly, EGFR-deficiency in hepatocytes also led to tumour nodules with increased size in the same model [6] which may be linked to less efficient DDR in the absence of EGFR signalling. This suggests that ADAM17-mediated release of EGFR ligands from myeloid cells is necessary to promote efficient DNA damage repair in hepatocytes and thus prevent genomically unstable hepatocytes.

Further studies, for example using inducible deficiency of ADAM17 in myeloid cells, or cell type-specific inhibition of ADAM17 are needed to determine whether myeloid ADAM17 is a potential drug target for the treatment of established malignant liver disease.

In conclusion, we have identified a dual role for ADAM17 in hepatic DNA damage repair and in the promotion of liver tumourigenesis.

## Acknowledgment

We thank Elena Voglas and Fabian Neumann for excellent technical assistance. We are grateful to the members of the Victor-Hensen animal facility at the Christian-Albrechts-University Kiel and to members of the central animal facility at the Paris-Lodron-University Salzburg.

## Author’s contribution

M.R., B.H., J.B., A.S., F.K., L.C., R.B. performed experiments, data analysis and contributed to writing of the manuscript. J.R., J.H.H., S.R.-J. contributed to the design of the study and writing of the manuscript. D.S.-A. designed the study, supervised and performed experiments, performed data analysis and wrote the manuscript.

