## Supplementary Material for "ADAM17 regulates hepatic DNA damage repair and tumour formation"

### Supplementary Experimental Procedures

Supplementary Table 1 – primary antibodies used in this study

| Target | Host species | Dilution | Distibuted by |
|---|---|---|---|
| **Western Blot** | | | |
| anti-ADAM10 (GTX63486) | rabbit | 1:1,000 | Genetex/Biozol, Eching Germany |
| anti-ADAM17 (ab39162) | rabbit | 1:1,000 | Abcam, Cambridge, UK |
| β-actin (A2066) | rabbit | 1:10,000 | Sigma-Aldrich, Steinheim, Germany |
| P-ATM (#4526) | mouse | 1:1,000 | Cell Signaling, Leiden, Netherlands |
| P-ATR (#2853) | rabbit | 1:1,000 | Cell Signaling, Leiden, Netherlands |
| cleaved Caspase 3 (#9661) | rabbit | 1:1,000 | Cell Signaling, Leiden, Netherlands |
| P-EGFR (GTX61353) | rabbit | 1:1,000 | Genetex/Biozol, Eching Germany |
| EGFR (#6627) | rabbit | 1:1,000 | Cell Signaling, Leiden, Netherlands |
| P-Erk1/2 (Thr202/Tyr204) (clone D13.14.4E) (#4370) | rabbit | 1:1,000 | Cell Signaling, Leiden, Netherlands |
| Erk1/2 (clone 137F5) (#4695) | mouse | 1:1,000 | Cell Signaling, Leiden, Netherlands |
| γH2Ax (#2577) | rabbit | 1:1,000 | Cell Signaling, Leiden, Netherlands |
| P-p38 (Thr180/Tyr182) (clone D3F9) (#4511) | rabbit | 1:1,000 | Cell Signaling, Leiden, Netherlands |
| p38 (clone D13E1) (#8690) | rabbit | 1:1,000 | Cell Signaling, Leiden, Netherlands |
| P-p53 (#9286) | mouse | 1:1,000 | Cell Signaling, Leiden, Netherlands |
| p53 (#2524) | mouse | 1:1,000 | Cell Signaling, Leiden, Netherlands |
| **Immunofluorescence** | | | |
| cleaved Caspase 3 (#9661) | rabbit | 1:1,000 | Cell Signaling, Leiden, Netherlands |
| CD11b (11-0112) | rat | 1:200 | eBiosciences/Thermo Fisher Scientific, Darmstadt, Germany |

| | | | |
|---|---|---|---|
| CLEC4F (AF2784) | goat | 1:250 | R&D Systems, Wiesbaden, Germany |
| P-EGFR (GTX61353) | rabbit | 1:250 | Genetex/Biozol, Eching Germany |
| F4/80 (MCA497R) | rat | 1:200 | Biorad, Feldkirchen, Germany |
| HNF4α (sc-6556) | goat | 1:400 | Santa Cruz, Heidelberg, Germany |
| γH2Ax (#2577) | rabbit | 1:200 | Cell signaling, Frankfurt a. M., Germany |
| ICAM-1 (#10-0542) | rat | 1:200 | eBioscience/Thermo Fisher Scientific, Darmstadt, Germany |
| In Situ Cell Death Detection Kit, Fluorescein (#11 684 795 910) | | | Roche, Rotkreuz, Switzerland |

Supplementary Table 2 – secondary antibodies used in this study

| Target | Host species | Dilution | Distibuted by |
|---|---|---|---|
| **Immunofluorescence** | | | |
| anti-rabbit AF488 (A11034) | goat | 1:200 | Thermo Fisher Scientific, Darmstadt, Germany |
| anti-rabbit AF594 (A21207) | donkey | 1:200 | Thermo Fisher Scientific, Darmstadt, Germany |
| anti-rabbit AF647 (A31573) | donkey | 1:200 | Thermo Fisher Scientific, Darmstadt, Germany |
| anti-rat AF488 A(21208) | donkey | 1:200 | Thermo Fisher Scientific, Darmstadt, Germany |
| anti-rat AF594 (A21209) | donkey | 1:200 | Thermo Fisher Scientific, Darmstadt, Germany |
| anti-rat AF650 (SA5-10029) | donkey | 1:200 | Thermo Fisher Scientific, Darmstadt, Germany |
| anti-goat AF594 (A11058) | donkey | 1:200 | Thermo Fisher Scientific, Darmstadt, Germany |
| anti-rabbit TSA Fluorescein (NEL701A001KT) | | | Perkin Elmer, Rodgau, Germany |
| anti-rabbit TSA (NEL744001KT) | | | Perkin Elmer, Rodgau, Germany |
| EnVision+ System HRP (K4003) | | | Dako/Agilent, Glostrup, Denmark |

### Supplementary Figures

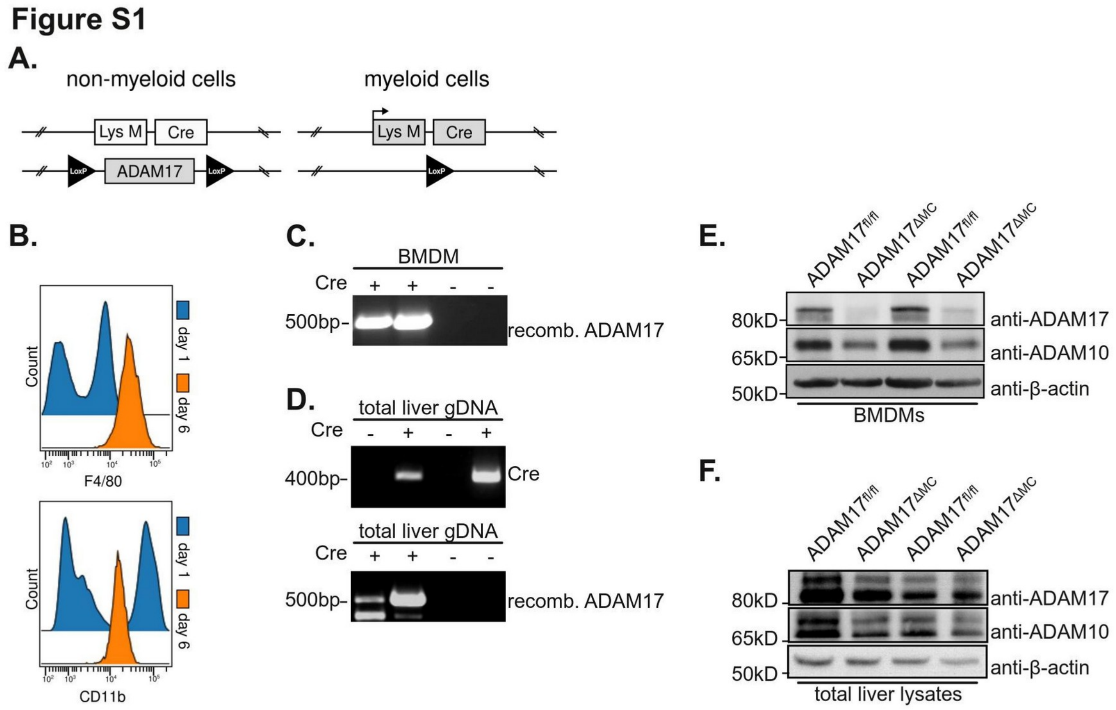


**Figure S1: Myeloid-specific deletion of ADAM17. A.** Recombination strategy. *Adam17* exon 2 is flanked by loxP sites and is excised in the presence of Cre recombinase. **B.** F4/80 and CD11b expression in cultures of bone-marrow derived macrophages (BMDMs) at day 1 and day 6. **C.** RT-PCR detection of Cre expression in BMDMs. D. PCR detection of Cre and recombined *Adam17* allele in total liver genomic DNA (gDNA). **E.** Immunoblot detection of ADAM17 and ADAM10 demonstrates efficient deletion of ADAM17 in BMDM derived from LysM-Cre x ADAM17$^{fl/fl}$ mice. **F.** Immunoblot detection of ADAM17 and ADAM10 expression in total liver lysates.

### Figure S2

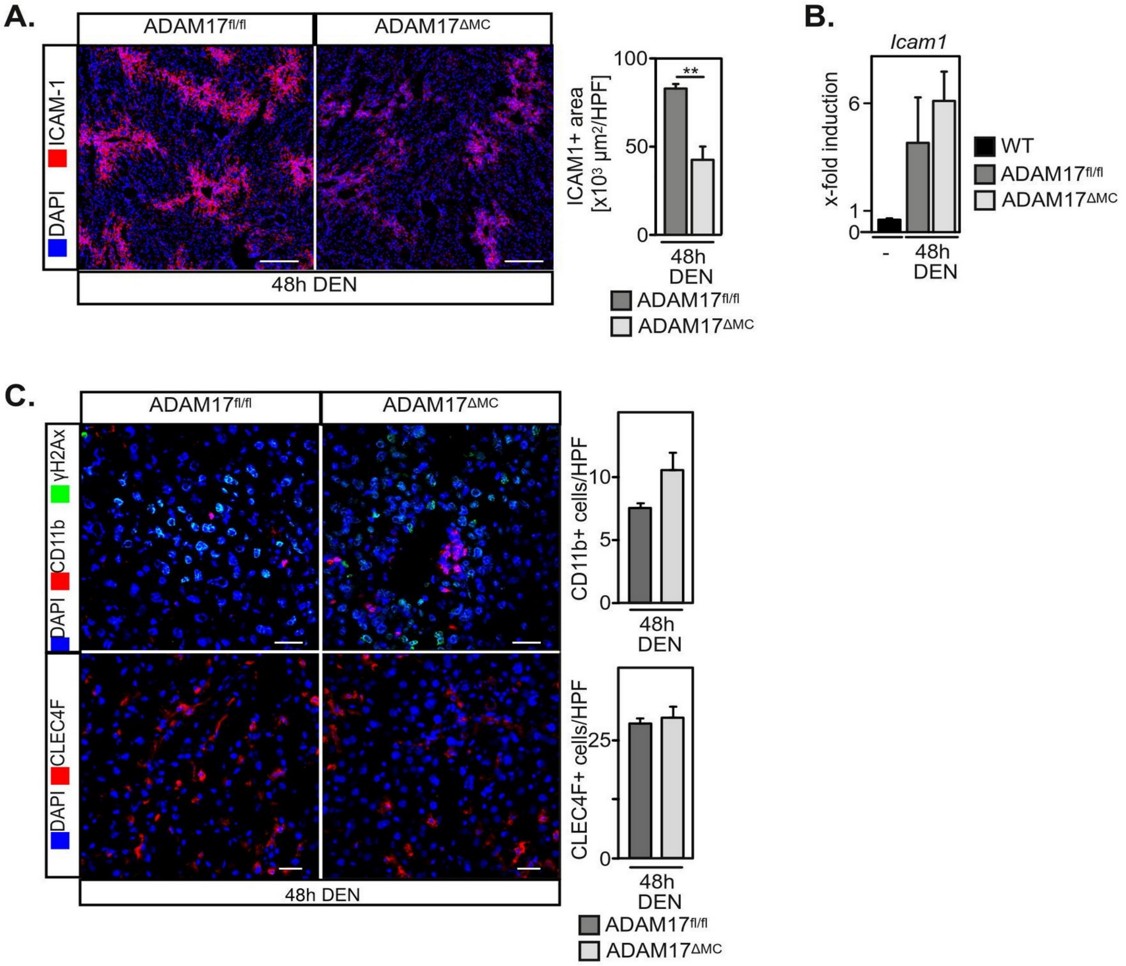


**Figure S2: A.** Immunofluorescence reveals pericentral expression of ICAM1 which appears to be reduced in mice with myeloid-specific ADAM17-deficiency. n=3-6 mice/group, ***P*<0.01 by Wilcoxon signed rank test. **B.** Expression of *Icam1* is increased upon acute DNA damage but unaltered in the absence of myeloid ADAM17. n=3 mice/group, Wilcoxon signed rank test. **C.** The number of infiltrating CD11b+ myeloid cells and resident CLEC4F+ Kupffer cells is unaltered in the absence of myeloid ADAM17. n=3-6 mice/group, unpaired two-tailed Student's *t* test.

Figure S3

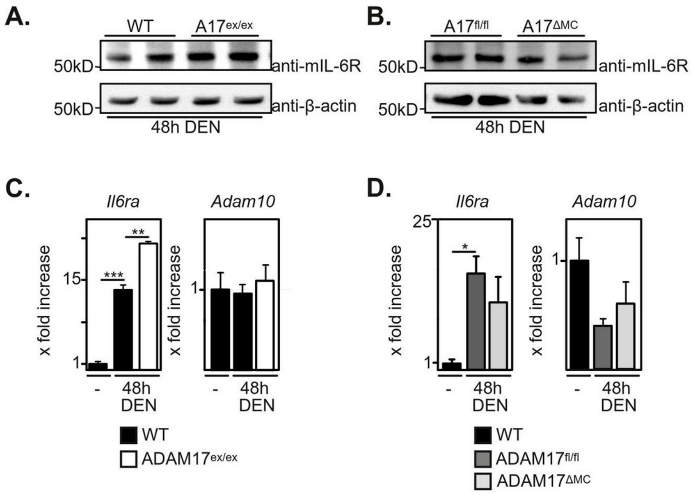


**Figure S3: A+B.** Protein levels of the IL-6 receptor (IL-6R) in total liver lysates is unaltered in the absence of ADAM17. **C+D.** Transcription of *Il6ra* is increased upon acute DNA damage and increased in the ubiquitous absence of ADAM17. Expression of *Adam10* is largely unaltered upon acute DNA damage and does not differ between control and ADAM17-deficient animals. n=3 mice/group, **P*<0.05, ***P*<0.01, ****P*<0.001, unpaired two-tailed Student's t test (C: *Il6ra, Adam10*; D: *Il6ra*), Wilcoxon signed rank test (D: *Adam10*)

**Figure S4**

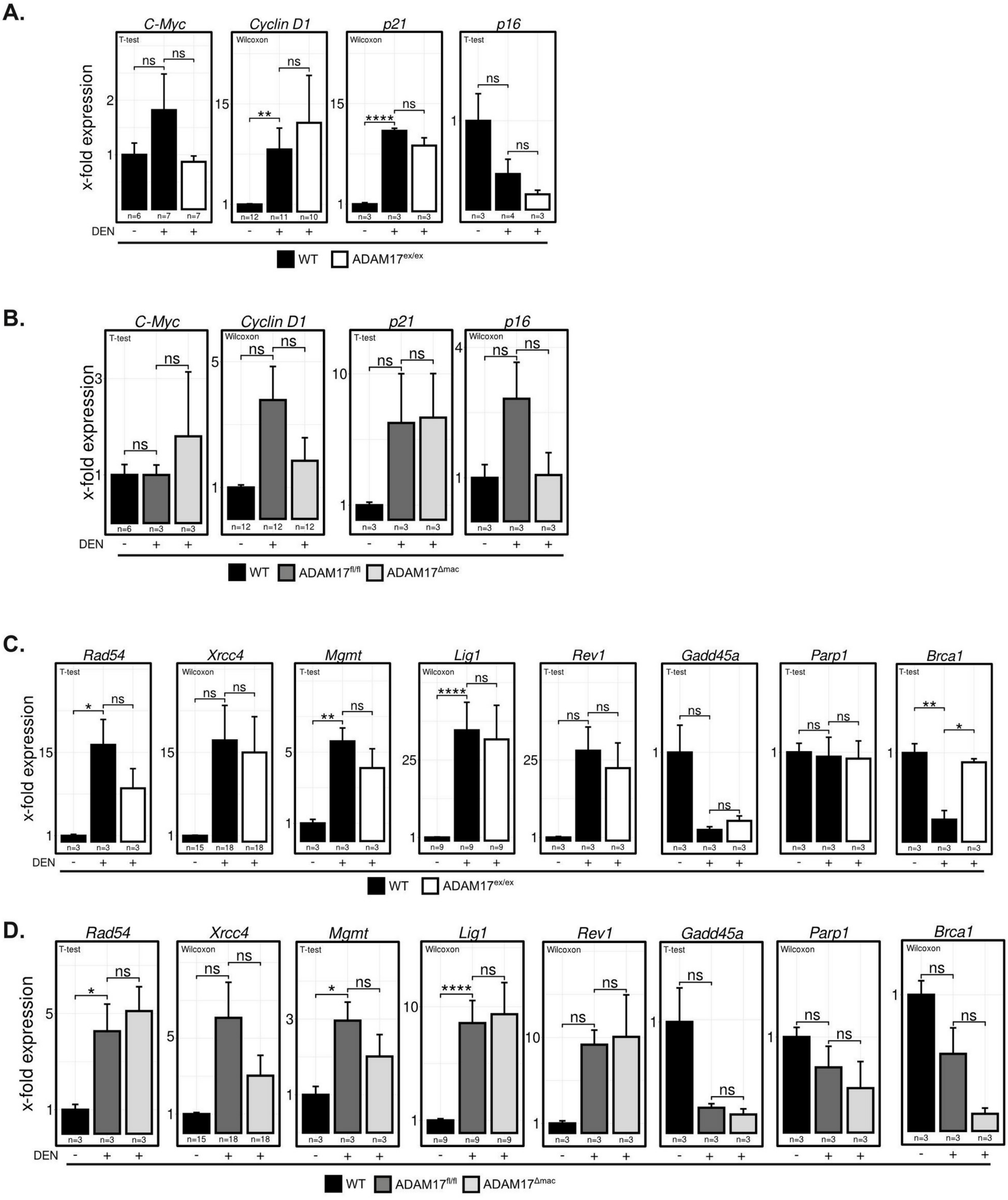


**Figure S4. A+B:** Transcription of genes that are involved in the cell cycle control is increased upon acute DNA damage but independent of genetic ADAM17-deficiency. n=1-4 mice/group, $^{**}P$<0.01, $^{***}P$<0.001, unpaired two-tailed Student's *t* test (A: *c-myc*, *Cyclin D1*, *p21, p16,;* B: *c-myc, p21*) or Wilcoxon signed rank test (B: *Cyclin D1, p16*) **C+D.** Transcription of genes involved in DNA damage repair is increased upon acute DNA damage but independent of genetic ADAM17-deficiency. n=1-4 mice/group $^{*}P$<0.05, $^{**}P$<0.01, $^{***}P$<0.001, unpaired two-tailed Student's *t* test (C:*Gadd45b, Brca1, Rad54,*

*Mgmt, Rev1, Gadd45a, Parp1;* D: *Rad54, Mgmt, Gadd45a*) or Wilcoxon signed rank test (C: *Xrcc4, Lig1*; D: *Mdm2, Brca1, Lig1, Xrcc4, Parp1, Rev1, Gadd45b*)
